## Supplemental Tables for "Oral rabies vaccination of dogs – experiences from a field trial in Namibia"

Supplementary table 1. The mean, minimum and maximum vaccination success rate (%) and the results of the univariate analysis of the selected independent variables (n = number of settings)

| Variable | n | mean | minimum | maximum | p-value (statistical test) |
|---|---|---|---|---|---|
| date | **8** | 82.5 | 69.8 | 89.0 | 0.0048 (Chi²) |
| period of the day | **4** | 82.5 | 81.0 | 84.5 | 0.8790 (Fisher) |
| team | **4** | 82.5 | 80.2 | 85.3 | 0.4263 (Chi²) |
| level of supervision 1) | **3** | 82.5 | 73.3 | 82.8 | 0.4044 (Chi²) |
| social status 2) | **2** | 82.6 | 80.8 | 85.9 | 0.0494 (Fisher) |
| Size 3) | **3** | 82.4 | 77.2 | 87.2 | 0.0166 (Chi²) |
| Sex 4) | **2** | 82.9 | 82.0 | 84.5 | 0.3700 (Fisher) |

1) 1 dog with no entry for level for supervision; not included in the statistical analysis'

2) 26 dogs with no entry for social status; not included in the statistical analysis

3) 14 dogs with no entry for size; not included in the statistical analysis

4) 30 dogs with no entry for sex; not included in the statistical analysis

Supplementary table 2: Odds ratios of the Multiple Logistic Regression (MLR) Model.

| coefficient | Variable | Odds ratios | 95% CI (profile likelihood) |
|---|---|---|---|
| β0 | Intercept | 0.4756 | 0.1819 to 1.110 |
| β1 | B[21.10.2021] | 0.6802 | 0.2724 to 1.870 |
| β2 | B[22.10.2021] | 0.7462 | 0.3063 to 2.016 |
| β3 | B[23.10.2021] | 0.4022 | 0.1543 to 1.139 |
| β4 | B[25.10.2021] | 0.3564 | 0.1257 to 1.065 |
| β5 | B[26.10.2021] | 0.4483 | 0.1568 to 1.350 |
| β6 | B[27.10.2021] | 0.5079 | 0.1893 to 1.467 |
| β7 | B[28.10.2021] | 1.125 | 0.4302 to 3.206 |
| β8 | C[multiple] | 0.6188 | 0.4203 to 0.8984 |
| β9 | D[large] | 1.254 | 0.6827 to 2.215 |
| β10 | D[small] | 0.5862 | 0.3905 to 0.8656 |

Supplementary table 3: The effect of bait handling (consumption - amount of bait matrix consumed; chewing time – total time [seconds] dog spend chewing on the bait; sachet – fate of sachet after bait consumption, discarded or swallowed) on vaccination success (%); N – number of dogs offered a bait, n – number of dogs considered successfully vaccinated. Data sets with an entry 'unknown' for these variables were excluded from statistical analysis.

| variable | N | n | % | p-value |
|---|---|---|---|---|
| <u>Consumption</u> | | | | |
| - complete (100%) | 731 | 682 | 92.3 | |
| - most (>50%) | 133 | 119 | 89.5 | 0.2822 (Chi²) |
| - little (<50%) | 10 | 9 | 90.0 | |
| <u>Chewing time</u> | | | | |
| - long (>60sec) | 89 | 76 | 85.4 | |
| - medium (30-60sec) | 318 | 297 | 93.4 | 0.0039 (Chi²) |
| - short (10-30sec) | 225 | 217 | 96.4 | |
| - very short (<10sec) | 242 | 219 | 90.5 | |
| <u>sachet</u> | | | | |
| - discarded | 425 | 384 | 90.4 | 0.0135 (Fisher) |
| - swallowed | 442 | 419 | 94.8 | |
